# Recording and Manipulation of the Maternal Oxytocin Neural Activities in Mice

**DOI:** 10.1101/2021.07.26.453888

**Authors:** Hiroko Yukinaga, Mitsue Hagihara, Kazuko Tsujimoto, Hsiao-Ling Chiang, Shigeki Kato, Kazuto Kobayashi, Kazunari Miyamichi

## Abstract

Pulsatile release of the hormone oxytocin (OT) mediates uterine contraction during parturition and milk ejection during lactation^1–3^. These pulses are generated by unique activity patterns of the central neuroendocrine OT neurons located in the paraventricular and supraoptic hypothalamus. Classical studies have characterized putative OT neurons by *in vivo* extracellular recording techniques in rats and rabbits under anesthesia^1, 4–7^ or awake^8–10^. Due to technical limitations, however, the identity of OT neurons in these previous studies was speculative based on their electrophysiological characteristics and axonal projection to the posterior pituitary, not on *OT* gene expression. To pinpoint OT neural activities among other hypothalamic neurons that project to the pituitary^11, 12^ and make better use of cell-type-specific neuroscience toolkits^13^, a mouse model needs to be developed for studies of parturition and lactation. We herein introduce viral genetic approaches in mice to characterize the maternal activities of OT neurons by fiber photometry. During lactation, a sharp photometric peak of OT neurons appeared at approximately 520 s following simultaneous suckling stimuli from three pups. The amplitude of the peaks increased as the mother mice experienced lactation, irrespective of the age of the pups, suggesting the intrinsic plasticity of maternal OT neurons. Based on a mono-synaptic input map to OT neurons, we pharmacogenetically activated the inhibitory neurons in the bed nucleus of the stria terminalis and found suppression of the activities of OT neurons. Collectively, our study illuminates temporal dynamics in the maternal neural activities of OT neurons and identifies one of its modulatory circuits.

**Highlights:**

- Pulsatile activities of genetically-defined OT neurons in mother mice were recorded *in vivo*.
- The maternal experience-dependent plasticity of the OT neural activities was found.
- Input-mapping of OT neurons in mother mice was performed by rabies-mediated trans-synaptic tracing.
- Photometric peaks of OT neurons were suppressed by the activation of BST inhibitory neurons.

## RESULTS

During parturition and lactation, OT neurons are thought to fire synchronously in bursts, resulting in pulsatile OT secretion, which is necessary for the contraction of the uterine and mammary glands^1, 2^. Although cell-type-specific recoding and manipulations of the OT system have been reported in previous studies of OT-mediated emotions and social behaviors^3, 14–18^, the maternal functions of OT neurons remain uncharacterized by cell-type-specific neuroscience toolkits. This motivated us to introduce *in vivo* chronic Ca^2+^ imaging to characterize the activities of OT neurons during parturition and lactation.

### Activities of OT neurons during parturition

We used fiber photometry^19^ to monitor selectively the activity dynamics of genetically-defined OT neurons in free-moving female mice. As a target, we focused on the paraventricular nucleus of the hypothalamus (PVH) because it contains a large number of OT neurons projecting to the posterior pituitary^12^ and can be accessed with minimum damage to the other hypothalamic regions. Hereafter, for simplicity, we refer to OT neurons located in the PVH as OT neurons. An optical fiber was implanted above the PVH of sexually naïve *OT-Cre*^20^*; Ai162*^21^ double heterozygous female mice. These mice were then crossed with stud male mice, and Ca^2+^ imaging was performed. Post-hoc histochemical analyses confirmed the fiber location (Figure 1A) and specificity of GCaMP6s expression (Figure 1B and 1C).

**Figure 1.**
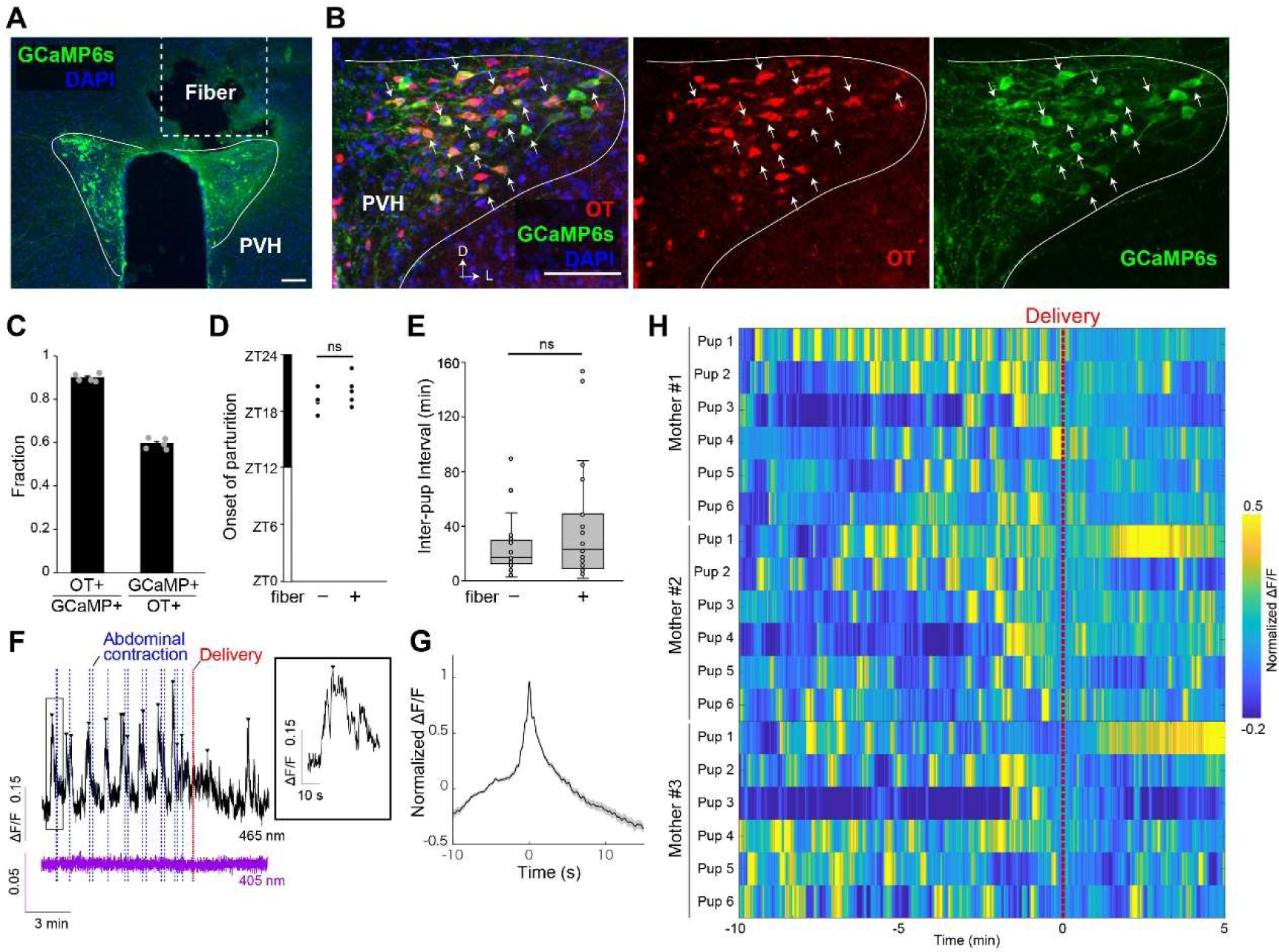
Fiber photometry recording of OT neurons during parturition. (A) Representative coronal brain section showing the location of the optical fiber and expression of GCaMP6s in the PVH. (B) Typical example of a 30-μm coronal section showing PVH stained by anti-OT (red) and anti-GFP (for GCaMP6s, green) antibodies counterstained with DAPI (blue). Arrows indicate some of the GCaMP6s-positive cells. Of note, most GCaMP6s-positive cells expressed OT, whereas a substantial number of OT neurons were GCaMP6s-negative. D, dorsal, L, lateral. (C) Quantification of specificity (OT+/GCaMP6s+) and efficiency (GcaMP6s+/OT+). *n* = 5 mice each. The efficiency of GCaMP6s expression was 59.0% ± 1.0%. This suboptimal efficiency might reflect the stochastic inactivation of the *tetracycline response element* promoter in the *Ai162* line. (D) Temporal distribution of the initiation of parturition under the fiber photometry setting (fiber +). No difference was found compared with control animals without connecting to the optical fiber (fiber –). ns, p > 0.05 by two-sided *t*-test. (E) Quantification of the pup delivery interval under the fiber photometry setting (fiber +). No difference was found compared with control animals without connecting to the optical fiber (fiber –). The center line shows the median; box limits represent upper and lower quartiles. ns, p > 0.05 by two-sided *t*-test. (F) Representative peri-event photometry traces of the 405-nm channel (non– calcium-dependent background, purple) and the 465-nm channel (calcium-dependent GCaMP6s signals, black) showing –10 min to +5 min relative to delivery. The photometric peaks are indicated by arrowheads, and the timing of abdominal contractions and pup delivery are represented by blue and red vertical dotted lines, respectively. Of note, this sample (mother #2-pup 2) corresponds to the data shown in Movie S1. The inset shows the high magnification view of the boxed area. (G) The mean of normalized peri-event traces of the peaks observed in mothers from – 10 min to delivery (*n* = 6 mothers, 292 peaks). The shadow represents the SEM. (H) Colored heat map representation of normalized ΔF/F from –10 min to +5 min relative to delivery for three representative mothers. Bin = 0.5 s. For each mother, the data of the first 5–6 deliveries are analyzed, because after the sixth delivery, the nest became too crowded to analyze the timing of the birth. We noticed a significant reduction of peaks after delivery: within the 5-min bin, the averaged peak number was 4.5 ± 0.44 (before) and 2.4 ± 0.38 (after) (p < 0.001 by two-sided *t*-test). Scale bar, 100 μm.

To monitor the neural activities of OT neurons during parturition, we applied an apparatus to capture the Ca^2+^ dynamics together with video records of the side and bottom views of the mouse cage (Movie S1). We habituated the female mice to the apparatus and started recording from the evening before the expected parturition day. Parturition in our experimental condition started mostly from Zeitgeber time (ZT) 17 to 20 (Figure 1D), where ZT 0 is the onset of the light phase. It lasted for one to a few hours, with a median interval of about 20 min for each pup delivery. No difference was found in the inter-pup interval with or without connecting to optical fibers (Figure 1E), suggesting that spontaneous parturition under fiber photometry was grossly normal.

Fiber photometry data showed pulsatile Ca^2+^ activities that started about 10 min before each delivery, whereas no peaks were observed in the 405-nm channel representing a non–calcium-dependent signal (Figures 1F, 1G, and S1A–S1D). From the side view, we observed abdominal contractions of the mother occurring 10–15 s after the photometric peaks (Figure 1F). The number of peaks before each delivery was often 6–8, but this varied substantially, even among individual deliveries of pups by a single mother (Figure S1A–S1D). In a heat map representation, high Ca^2+^ signals were clustered before, but not after, the delivery of pups (Figure 1H), while some intensive signals after the delivery of pups likely reflected the delivery of placental materials. These data show the temporal dynamics of photometric signals of OT neurons in the course of spontaneous parturition in mice.

### Activities of OT neurons during lactation

We recorded the activities of OT neurons of female mice in various conditions. While no apparent photometric peaks were observed when late pregnant female mice played with a toy (Figure S1E), we detected intensive photometric peaks during lactation starting several hours after parturition (Figure S1F). These observations motivated us to characterize further the neural activities of OT neurons in lactating mothers. A representative trace in Figure 2A shows photometry data from a mother at postpartum day 12 (PPD12) for 12 continuous hours. OT neurons often exhibited a cluster of peaks throughout the recording session. In this trace, we detected 62 peaks, with highly stereotyped height and full-width at half-maximum (FWHM) of individual waveforms (Figure 2B–2D). By contrast, the inter-peak interval substantially varied; although the majority clustered around 270 s, small fractions ranged widely, from 20 min to close to 1 h (Figure 2E).

**Figure 2.**
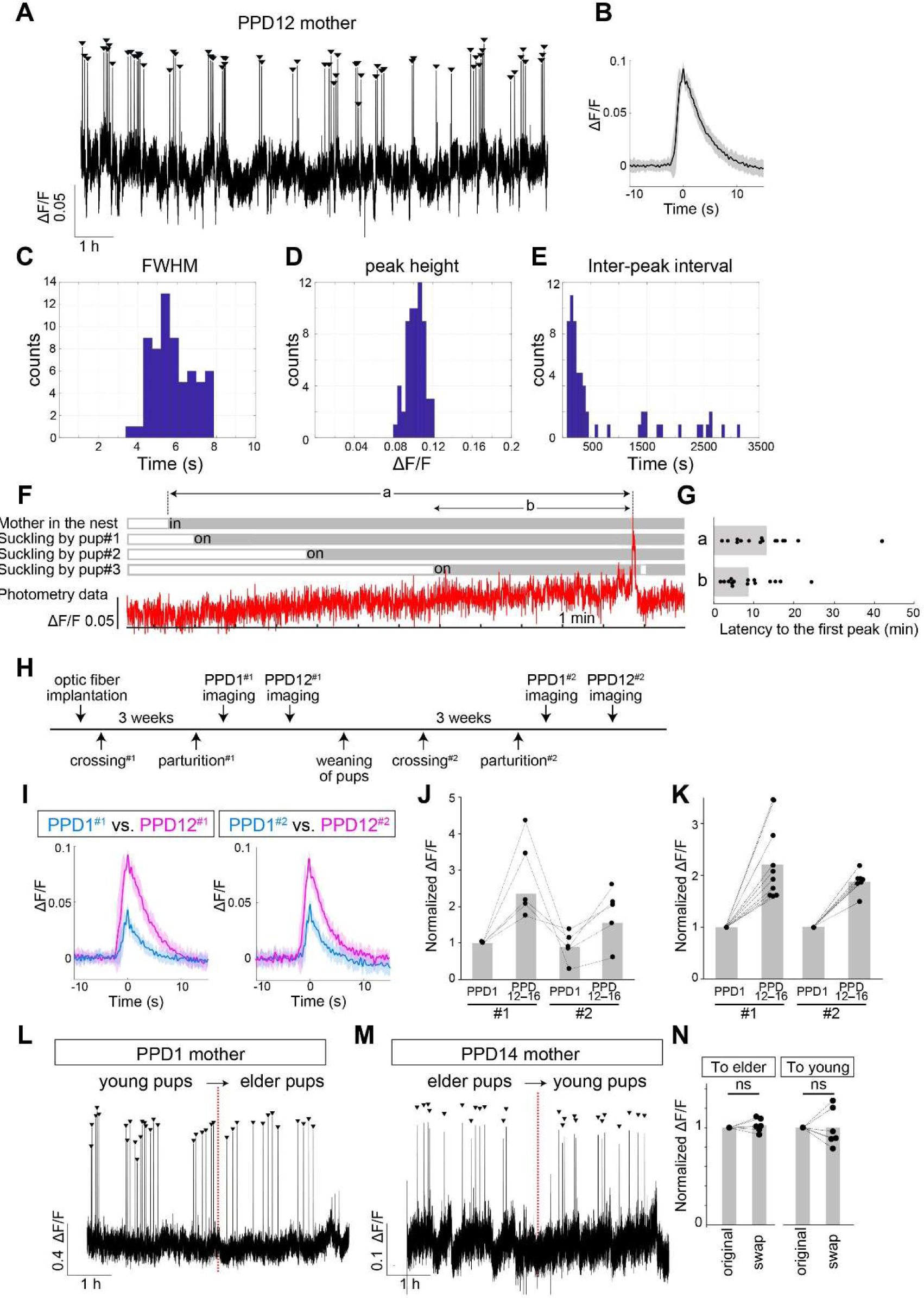
Fiber photometry recording of OT neurons during lactation. (A) Representative 12-h continuous fiber photometry trace from a PPD12 mother containing 62 photometric peaks. (B) Averaged peri-event trace of these peaks, with shadows showing the standard deviation (SD). (C–E) Histograms showing the FWHM (C), peak height (D), and inter-peak interval (E) from the raw data shown in (A). (F) Behavioral analysis of a mother and pups before each photometric peak. Three pups were placed with the PPD10 mother. Top, raster plot showing a mother staying in the nest (gray) and pups demonstrating suckling behaviors (gray). Bottom, photometric signals of the OT neurons. (G) Quantification of the latencies to the photometric peak from the mother returning to the nest (a) or from the third pup starting the suckling behaviors (b). *n* = 15 cases observed with *n* = 3 mothers. (H) Timeline of the experiments. #1 indicates the first round of parturition to lactation, whereas #2 indicates the second round after weaning and re-crossing. (I) Representative examples of averaged peri-event traces of individual peaks from a mother at four time points: PPD1 (blue) and PPD12 (magenta) of the #1 and #2 lactation. (J) The peak height normalized to the mean of that of PPD1^#1^ for *n* = 5 mothers in whom we could monitor OT neurons from PPD1^#1^ to PPD12–16^#2^. A significant difference was not supported by repeated measures one-way ANOVA with post hoc paired t-test with Bonferroni correction. (K) Peak height was calculated for each animal and shown as the fold change from PPD1^#1^ to PPD12–16^#1^ or PPD1^#2^ to PPD12–16^#2^ of the same animal. **, p < 0.01 and *, p < 0.05 by two-sided *t*-test with the Bonferroni correction. *n* = 10 for #1 and *n* = 7 for #2 data. (L) Representative examples of 5.5-h traces of fiber photometry data obtained from PPD1 (L) or PPD14 (M) mothers during lactation. In the middle of the recording session, the original pups were removed and foster pups were introduced. Arrowheads represent the photometric peaks. (N) Quantification of the fold change of peak height before (original) and after swapping pups. The p-values were 0.36 and 0.45 by two-sided *t*-test for PPD1 mothers (*n* = 6) and PPD12–16 mothers (*n* = 6), respectively.

Next, we aimed to characterize the relationships between nipple stimulation by pups and photometric signals. Based on the finding that three or more pups are required for reliable activation of a mother’s OT neurons (Figure S1G and S1H), we placed three pups with PPD10 mothers and monitored the behaviors of mothers and pups at the onset of photometric peaks (Movie S2). We found that the peak appeared on average at 520 ± 101 s (mean ± standard error of the mean) after simultaneous suckling by all three pups (Figure 2F and 2G). This supports the notion that the observed photometric peaks are related to milk ejection.

When we chronically monitored the activities of OT neurons in the early and later lactation periods, we noticed that individual peaks became much higher as mothers experienced lactation. Because the intensities of the photometric peaks observed in each mother varied considerably, probably because of the variable optical fiber location relative to the PVH, the exact value of ΔF/F could not be compared among different animals. Therefore, we analyzed the fold change of the same animal from PPD1 to PPD12–16 and found that the height of the peaks was almost doubled in the PPD12–16 compared with the PPD1 mothers (Figure 2H–2K). Enhanced photometric signals in the experienced mothers may have been caused by stronger nipple sucking by elder pups. Alternatively, this change may be intrinsic to maternal OT systems, independent of the sucking skills of pups. To distinguish these two possibilities, we cross-fostered pups between PPD1 and PPD12–16 mothers. The height of the peaks would change after swapping if the pups were the determinant. We found that the peaks in PPD1 and PPD12–16 mothers remained unchanged, regardless of the age of the pups (Figure 2L– 2N). Therefore, enhanced photometric peaks in PPD12–16 mothers are not simply due to the greater nutritional requests by pups, which suggests the presence of plasticity in the OT neural system. This can lead to a more efficient milk ejection reflex as pups grow.

If the photometric peaks were enhanced by lactation experience, how would they behave in secundiparous mothers who nurse their second litters after weaning the first ones? In some cases (*n* = 5), we chronically monitored the same female mice throughout the first (#1) and second (#2) postpartum periods (Figure 2I). Enhanced peaks established by PPD12^#1^ tended to return to baseline at PPD1^#2^ and increased again by PPD12^#2^ (Figure 2J); however, due to the limited numbers of samples, we could not confirm this notion statistically. Instead, we confirmed a significant increase in the peak height analyzed separately by the fold change from PPD1^#2^ to PPD12–16^#2^ (*n* = 7 secundiparous mothers, Figure 2K). These observations are consistent with the notion that the lactation-dependent enhancement of photometric peaks of OT neurons is reset by each parturition. Collectively, our data reveal the temporal dynamics of neural activities of OT neurons during lactation.

Lastly, we compared the waveforms of photometric signals during parturition (Figure 1G) and lactation (Figure 2B) using the same female mice. From parturition to PPD1, the average FWHM was significantly reduced and the average peak height was greatly increased (Figure S1I and S1J). Therefore, the photometric signals in lactation were steeper. The inter-peak interval was much shorter during parturition; the median peak interval was 50.3 s before delivery (Figure S1D) and about 270 s during lactation (Figure 2E). These data reveal that the shape and interval of photometric signals significantly differed between parturition and lactation, suggesting that the neural circuit mechanisms that generate these peaks change quickly (within several hours) from the peri-partition to lactation phase (Figure S1F).

### Input neural circuit mapping to OT neurons

How are the neural activities of OT neurons modulated by afferent neural circuitry? The current understanding is mostly based on the classical non–cell-type specific lesion or electric stimulation data in postpartum rats during lactation^22–24^. Mapping mono-synaptic input to OT neurons would be the first step to address this issue. We applied Cre-dependent rabies-virus-mediated trans-synaptic tracing^25, 26^ to OT neurons. To map long-distance input to OT neurons efficiently, we injected Cre-dependent adeno-associated virus (AAV) vectors for *CAG-FLEx-TC^b^*(*TVA-mCherry^bright^*) and *CAG*-*FLEx-RG* (rabies glycoprotein) into the PVH of sexually naïve *OT-Cre* female mice. We then crossed them with stud male mice and injected rabies *dG-GFP*+EnvA into the PVH of age-matched sexually-experienced nonpregnant control mice and PPD1 mothers (Figure 3A) for a side-by-side comparison. A negative control experiment omitting Cre expression showed 385.8 ± 76.4 GFP expressing (GFP+) cells as nonspecific rabies labeling^26^, the vast majority of which were located in the PVH (72%) and paraventricular thalamus (PVT, 18%). Thus, we excluded PVH and PVT from the analysis of long-range input to OT neurons.

**Figure 3.**
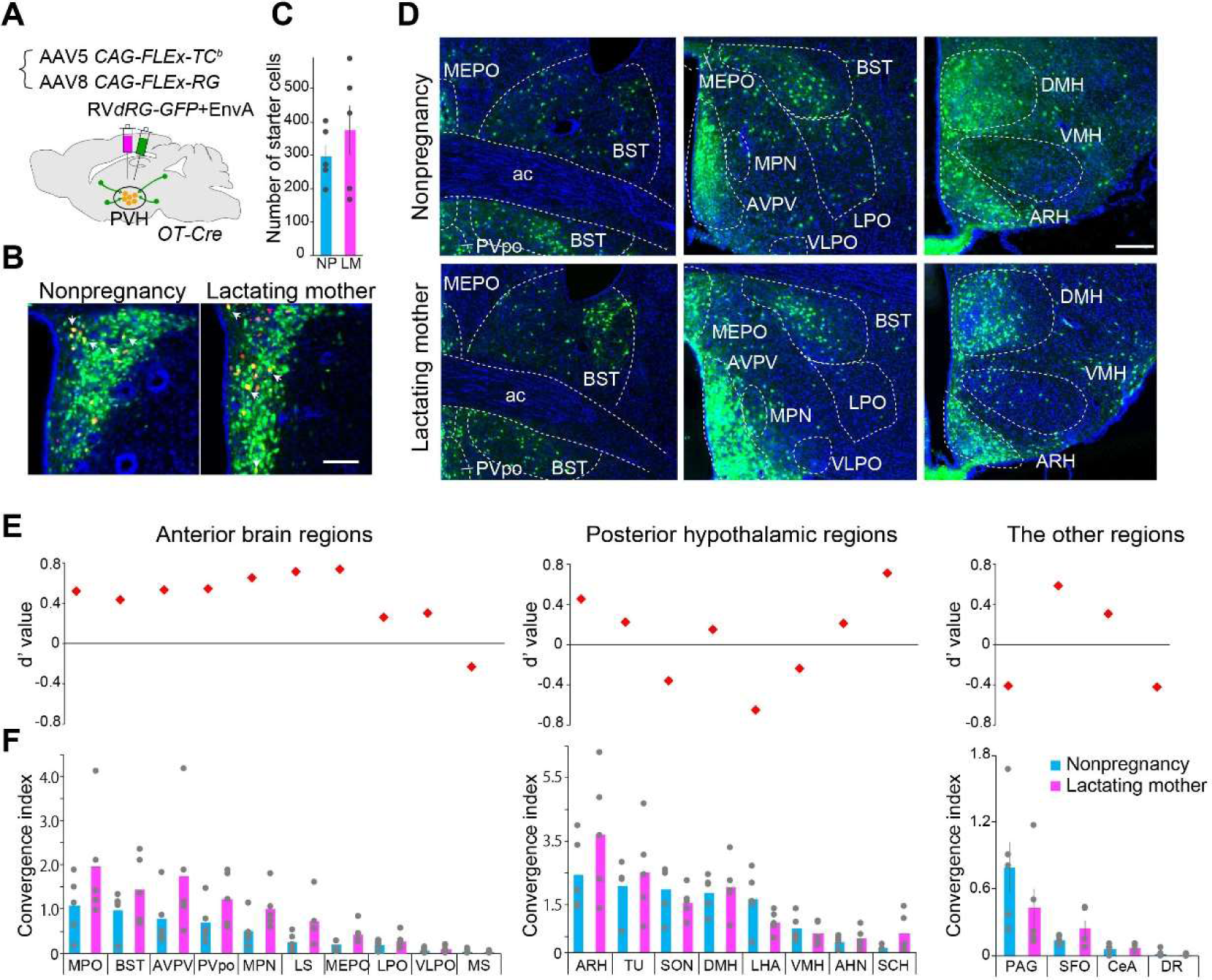
Quantification of input cells to OT neurons in the postpartum female mice. (A) Schematics of the experimental design. (B) Coronal sections of *OT-Cre* tracing brains showing the starter cells (defined by co-expression of TVA-mCherry and rabies-GFP, some indicated by arrows) in the PVH. Scale bar, 100 μm. (C) Quantification of starter cells per animal. No significant difference was found between nonpregnancy (NP) and lactating mothers (LM) by the Wilcoxon rank-sum test (p = 0.48). *n* = 5 each. (D) Representative coronal sections showing the distribution of presynaptic partners. Scale bar, 200 μm. (E, F) The d’-values and average convergence index (defined by the number of rabies-GFP+ cells normalized to the number of starter cells) with individual animal data in gray. *n* = 5 each. For abbreviations of brain regions, see Table S2.

Overall, the number of starter cells (i.e., those labeled with TVA-mCherry and rabies-GFP) was comparable in nonpregnant and lactating female mice (Figure 3B and 3C), consistent with a previous report showing no major changes in the number of OT neurons upon lactation in rats^17^. The majority of presynaptic neurons were located in the broad preoptic and hypothalamic areas (Figure 3D) with prominent input structures, including the medial preoptic nucleus (MPN), supraoptic nucleus (SO), dorsomedial hypothalamus (DMH), ventromedial hypothalamus (VMH), and arcuate hypothalamic nucleus (ARH). We also noticed substantial input cells in the extra-hypothalamic areas, such as the lateral septum (LS), the bed nucleus of the stria terminalis (BST), and the periaqueductal gray of the midbrain (PAG). Input patterns were grossly similar between nonpregnant female mice and lactating mothers (Figure 3E and 3F), with no statistically significant differences in any brain region (Table S1); however, a weak tendency to send more input to OT neurons was seen in anterior brain regions in lactating mothers.

Next, we analyzed the cell type of input neurons. Brain slices containing six selected areas were stained by *in situ* hybridization (ISH) with an excitatory or inhibitory neural marker, *vGluT2* or *vGAT*, together with immunostaining of rabies-GFP (Figure S2). Input neurons originating from the anteroventral periventricular nucleus (AVPV) and DMH were mixed with *vGluT2*+ and *vGAT*+ subpopulations, whereas those from the LS, BST, and ARH were mostly GABAergic, and those from the VMH were mostly glutamatergic. No difference in the fraction of excitatory or inhibitory neurons was observed between nonpregnant female mice and lactating mothers.

As TC^b^-based trans-synaptic tracing contained substantial local background labeling, we utilized a mutant TVA receptor^26^ to characterize local connectivity within the PVH more accurately and reduce the nonspecific labeling to zero (Figure S3). Rabies-GFP spread to local PVH neurons (Figure S3B), showing abundant local PVH input to OT neurons. To characterize the cell type of GFP-labeled neurons, we detected mRNA for OT and its related peptide hormone vasotocin^27^ (VT; also known as arginine-vasopressin), together with immunostaining of rabies-GFP (Figure S3D). In nonpregnant female mice, 6.2% ± 1.0% of GFP+ neurons were *OT+*, and 17.9% ± 2.4% were *VT+*, demonstrating intensive connectivity among neurons producing posterior pituitary hormones. In lactating females, OT-to-OT connection tended to increase, whereas no change was seen in VT-to-OT connection (Figure S3E). As VT and OT neurons account for only 20%–30% of GFP+ neurons in the PVH, other types of neurons are likely to contribute to the regulation of OT neurons via the local circuits.

In summary, our input circuit mapping forms a basis to characterize afferent neural circuitry to OT neurons that underlie various OT-mediated biological processes, including parturition, lactation, and social bonding^3^.

### Pharmacogenetic manipulations of maternal neural activities of OT neurons

The mono-synaptic input map to the OT neurons (Figure 3) can facilitate the study of circuit mechanisms by which maternal neural activities of OT neurons (Figures 1 and 2) are modulated by afferent circuitry. As a proof-of-principle, we combined *OT-Cre*-mediated fiber photometry recordings with cell-type-specific pharmacogenetic manipulations of a defined presynaptic structure. We focused on the inhibitory neurons in the BST (Figure 3D–3F, Figure S3) because they are the prominent long-distance input to OT neurons. To target inhibitory neurons selectively, we utilized a small enhancer sequence of the human *Dlx5*/6 intergenic region^28^ to drive hM3Dq or hM4Di for activation or inactivation of the targeted neurons^29^ (Figure 4A), with a negative control driving only mCherry. Post-hoc histochemical analyses revealed that the majority (94% ± 1.0%) of the transduced neurons were inhibitory based on the labeling with anti-GABA antibodies (Figure S4A) and that, on average, 85% of hM3Dq-dTomato+ and 80% of hM4Di-dTomato+ cells were located within the BST (Figure 4B, Figure S4B). For some data, we utilized AAV9 *CAG-FLEx-GCaMP6s* injected into the PVH instead of *Ai162* based on the confirmation that GCaMP6s was selectively expressed by OT neurons by using this method (Figure S4C and S4D).

**Figure 4.**
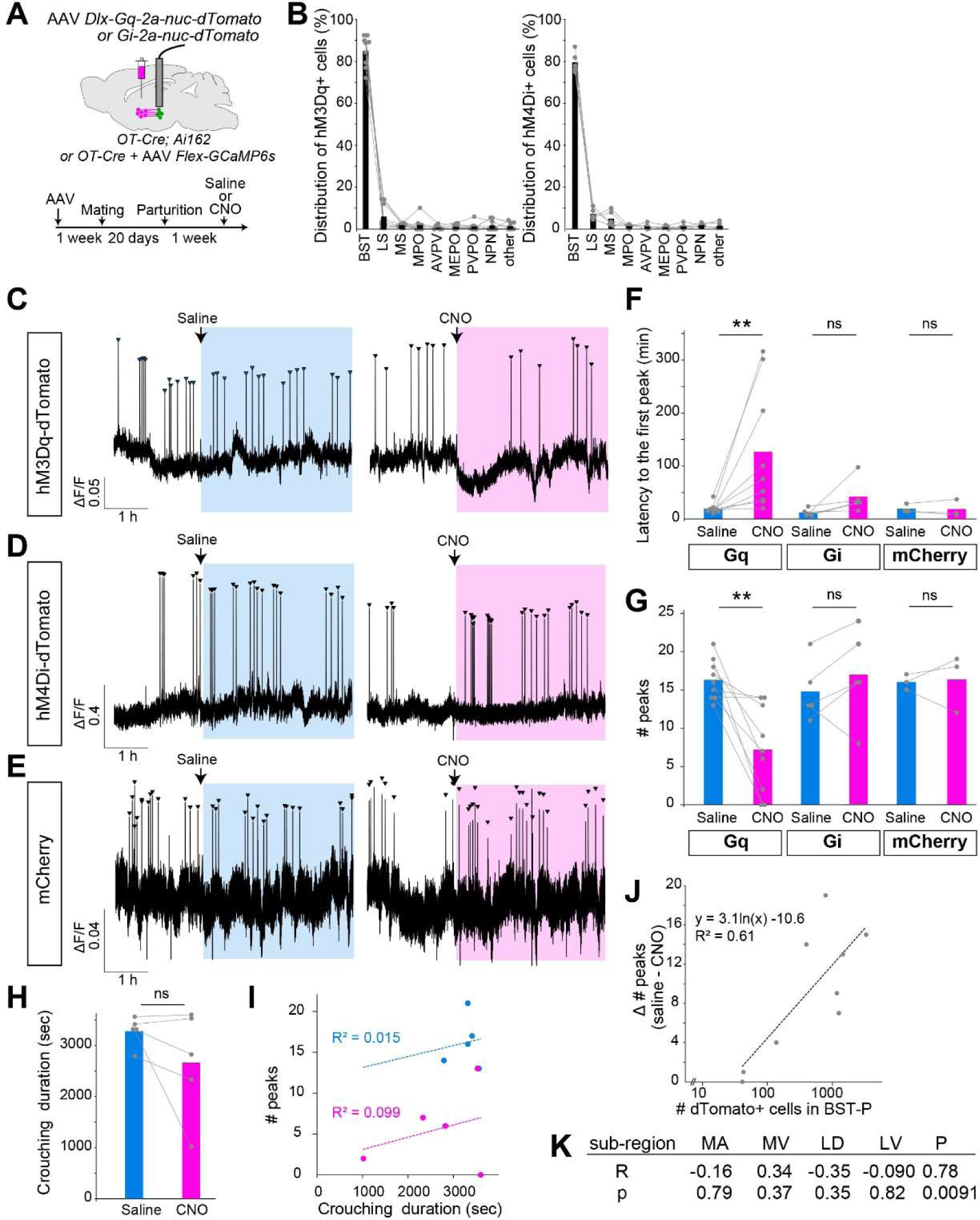
Modulation of the peak frequency by pharmacogenetic manipulations. (A) Schematics of the experimental design and timeline of the experiments. (B) The number of dTomato+ cells in each brain region normalized to the total number in all the regions, showing the distributions of hM3Dq+ or hM4Di+ neurons in the BST and surrounding regions. *n* = 9 for hM3Dq and *n* = 5 for hM4Di. For abbreviations of brain regions, see Table S2. (C–E) Representative 5.5-h traces showing the photometric peaks of OT neurons (arrowheads) in hM3Dq+ (C), hM4Di+ (D), or mCherry+ (E) mothers. The timing of saline or CNO injection is indicated by the vertical arrows. (F, G) Quantification of latency to the first peak (F) and the number of peaks in 3.5 h (G) after saline or CNO injection. *n* = 9 for hM3Dq, *n* = 5 for hM4Di, and *n* = 3 for mCherry control. **, p < 0.01 by two-sided Wilcoxon rank-sum test with the Bonferroni correction. (H) Crouching durations of hM3Dq+ mothers in a 1-h time window starting from 30 min after saline or CNO injection. No difference was found by the two-sided Wilcoxon rank-sum test. *n* = 5 mothers. (I) Correlation between crouching duration and the number of photometric peaks during saline (blue) or CNO (red) injected trials. No strong correlation was found. (J) Correlation of the number of dTomato+ cells (log scale) in the BST posterior (P) subdivision of hM3Dq+ mothers and the difference between the numbers of photometric peaks of saline and CNO trials. *n* = 9 mothers. (K) Correlation coefficient (R) and p-values in the analysis shown in (J) for five BST subdivisions. MA, medial anterior; MV, medial ventral; LD, lateral dorsal; LV, lateral ventral; P, posterior.

We performed a 6-h photometry recording twice in PPD6–13 mothers. On the first day, either saline control or clozapine-N-oxide (CNO) was intraperitoneally injected 2 h after the onset of the imaging session. On the second day, we replaced saline and CNO for counterbalancing. In the hM3Dq+ female mice, latency to the first peak was significantly elongated, and the number of peaks per 3.5-h time window was significantly decreased following CNO injection (Figure 4C, 4F, and 4G). No effect was observed in negative controls such as saline-injected trials or CNO injection into mCherry+ animals (Figure 4E). For some hM3Dq+ animals, we analyzed the crouching durations of mothers in the nest as an inference of maternal motivation. CNO injection did not significantly alter the crouching durations (Figure 4H). No correlation was found between the number of photometric peaks and the crouching duration (Figure 4I), suggesting that the hM3Dq-mediated effect is not primarily due to the loss of maternal interactions with pups. As the hM3Dq-mediated reduction of the photometric peaks varied substantially among individuals (Figure 4F), we next analyzed distributions of hM3Dq+ neurons within the subdivisions of the BST (Figure S4B). A positive correlation was found between the number of hM3Dq+ neurons in the BST posterior subdivision and the CNO-mediated reduction of the peak (Figure 4J), whereas no such correlation was found in the other subdivisions of the BST (Figure S4K). By contrast, in the hM4Di+ female mice, CNO injection did not affect the frequency of photometric peaks (Figure 4D, 4F, and 4G), despite it affecting the expression of *c-Fos*, a neural activity marker gene, in the local BST neurons (Figure S4E–4G). These data suggest that the inhibitory neurons in the BST, particularly those in its posterior subdivision, can modulate the neural activities of OT neurons during lactation; however, whether this effect is mediated by the direct presynaptic partners of OT neurons in the BST remains an open question.

## DISCUSSION

The present study expands the scope of cell-type-specific recording of OT neural activities^14, 15, 18, 30^ to maternal functions during parturition and lactation. Our fiber photometry data generally support single-unit recording of putative OT neurons in rats^1^. For example, peaks were initiated about 10 min before the delivery of the first pup (Figure 1), which is in-line with data from parturient rats^31^. The inter-peak interval of the photometric peaks during lactation was about 270 s (Figure 2E), close to the 300 s described in rats^4, 7^. The peaks started approximately 520 s after simultaneous suckling by three pups, also supporting the notion that the observed peaks are related to the milk ejection reflex. Notably, one limitation of our study is the lack of characterization of spiking or bursting activities of OT neurons required for OT release. In addition, we did not directly detect the contraction of the uterus/mammary gland mediated by the released OT. Future studies should attempt to link photometric signals directly with the action potentials and downstream endocrinological functions of OT.

We found the enhancement of individual photometric peaks from PPD1 to PPD12–16 mothers (Figure 2). As the age of pups did not significantly affect the intensity of the peaks, we speculate that this plasticity is mostly autonomous to the OT system of mothers. Of note, while devising the present study, a similar observation was reported in lactating rats^32^. What are the underlying mechanisms? In one scenario, the number of OT neurons forming each pulse is not changed from PPD1 to PPD12–16, but each neuron fires more. Alternatively, new OT neurons are recruited as mothers experience lactation. To distinguish these two possibilities, studies involving single-cell resolution imaging of OT neural activities are needed. We also found that the height, inter-peak interval, and waveform of the neural activities of OT neurons differed substantially between parturient and lactating females; therefore, it would also be interesting to investigate whether the OT neurons that are active during parturition also work for the milk ejection reflex, or whether distinct sub-populations of OT neurons exist for delivery and lactation.

The present study identified direct presynaptic partners of OT neurons by using a rabies-virus-mediated trans-synaptic tracing technique^25, 26^. Although a few other studies have reported similar tracing data in rats and mice in different biological contexts^15, 30, 33–35^, our data expand the scope of circuit analysis to lactating mothers and provide quantitative information on presynaptic structures and cell types (Figure 3 and Figure S2). In addition, we demonstrated the presence of local OT-to-OT and VT-to-OT connections (Figure S3). As the morphological changes in glia-OT neuron associations during lactation have been well documented^36^, we initially expected a drastic reorganization in input to OT neurons in mothers compared with nonpregnant female mice. However, our data suggest widespread and nuanced adjustments of connections (Figure 3 and Figure S3).

Nevertheless, our input map to OT neurons is a useful resource to characterize the afferent circuitry of OT neurons. To show this utility by pharmacogenetic manipulations, we identified a modulatory function of inhibitory neurons in the BST, particularly in its posterior subdivision (Figure 4). Our data not only confirm classical lesion and electronic stimulation data in rats^24, 37^, but also expand the scope to cell-type-specific and reversible manipulations of the neural activities of OT neurons in mothers. As diverse signals such as vomeronasal input^38^, stress, and anxiety^39^ are mediated by the BST, it would be interesting to ask whether BST input to OT neurons is indeed involved in the acute physical and/or mental stress-induced impairment of the milk ejection reflex, a phenomenon often reported in postpartum women^40^. More generally, our data form the basis for comprehensive circuit characterizations of various input nodes. A major obstacle in this direction is the necessity of Cre-based double transgenic mice for use in our method, which would not only limit the throughput of the data collection, but also compromise the use of various Cre-dependent toolkits for dissecting molecular and neural circuit functions. To overcome this issue, a simple Cre-free method for monitoring OT neural activities^32^ should be applied to mice. We expect that viral genetic manipulations of upstream neural activities or gene functions will illuminate the circuit mechanisms that shape the pulsatile activities of OT neurons.

## Supplementary Figures

**Figure S1.**
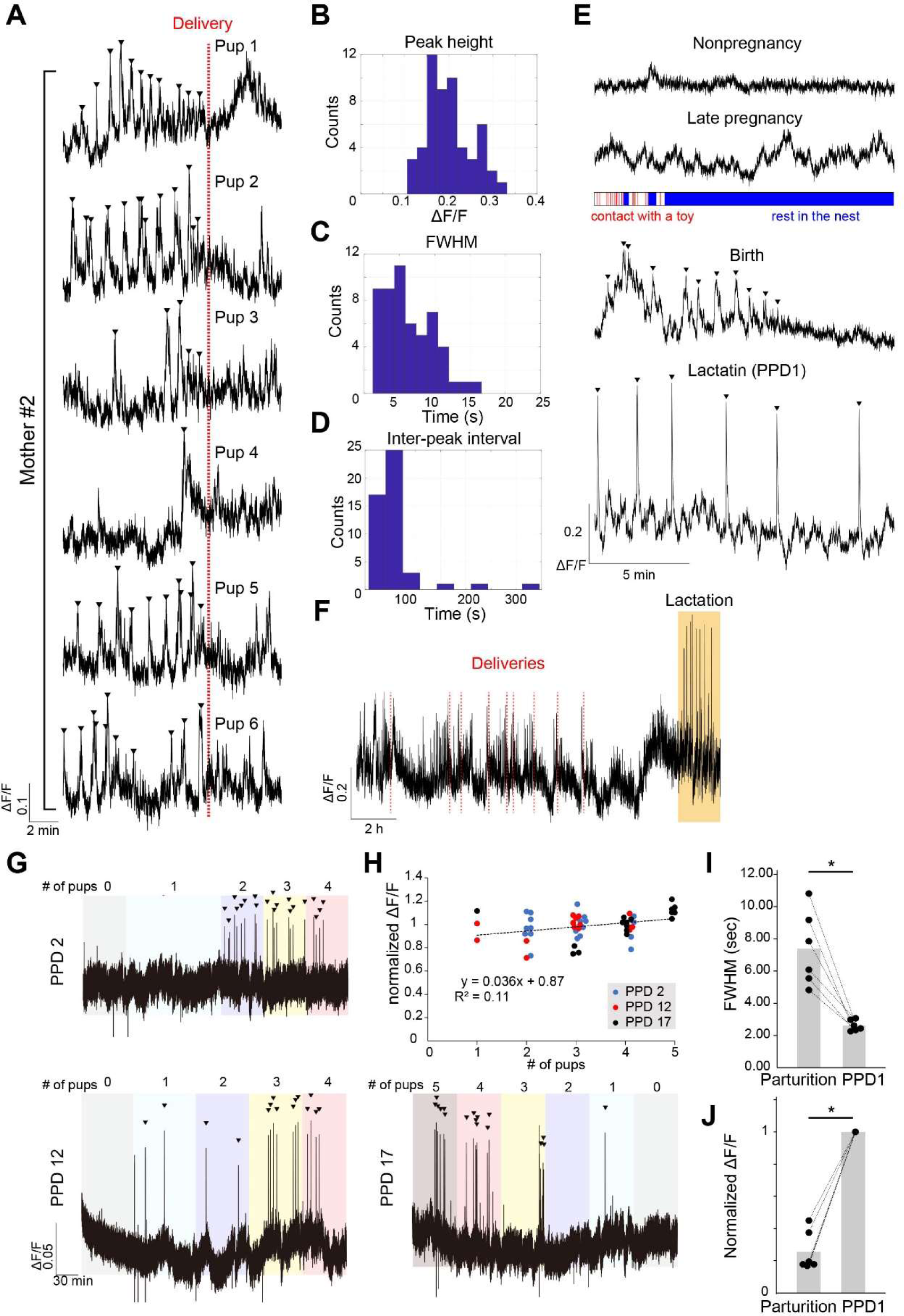
Photometric peaks during parturition in various reproductive conditions and with different numbers of pups, related to Figures 1 and 2. (A) Peri-event photometry traces from –10 min to +5 min relative to the delivery (indicated by the red vertical line) of each pup from a single female mouse (corresponding to mother #2 in Figure 1). The peaks detected before each delivery are shown by arrowheads. Of note, the clustered photometric peaks of OT neurons were not caused by the suckling of newborns, as we did not detect any suckling until the end of the delivery. (B–D) Histograms showing peak height (B), full width at half-maximum (FWHM, C), and inter-peak interval (D). Peaks that occurred from –10 min to delivery from the raw data shown in (A) were used for the calculation. (E) Representative examples of individual 15-min traces of photometry data of a female mouse obtained from different conditions as indicated. Nonpregnancy: a nonpregnant female mouse staying only in the home cage. Late pregnancy: the same mouse at gestation day 17 playing with a toy. The bottom raster plot shows the timing of contact with a toy in red. Birth: during partition. Lactation: during breastfeeding at postpartum day 1 (PPD1). Arrowheads indicate photometric peaks. (F) A representative example of photometry data showing activities of OT neurons from parturition to lactation. Deliveries of pups are indicated by the red vertical lines. About 4 h after the delivery of the last pup, the mother stably showed crouching behaviors, and intensive photometric peaks emerged, suggesting the beginning of the lactation period. Notably, the individual photometric peaks were higher during lactation compared with parturition. (G) Three representative examples of photometric peaks during lactation at different postpartum days (PPD2, 12, or 17) when the number of pups was changed from 0 to 5. Zero or one pup did not efficiently evoke the photometric peaks; two to three pups were necessary for the stable peaks (arrowheads). (H) Correlation between the number of pups and the normalized peak height. For each condition, the peak height was normalized to the mean of the peaks during lactation for four pups. A weak positive correlation was found (R = 0.33, p = 0.017). (I, J) Comparison of photometric signals of OT neurons during parturition and lactation. Average FWHM (I) and the fold change of peak height during parturition normalized to that of PPD1 (J). *, p < 0.05 by two-sided Wilcoxon rank-sum test. *n* = 6 each.

**Figure S2.**
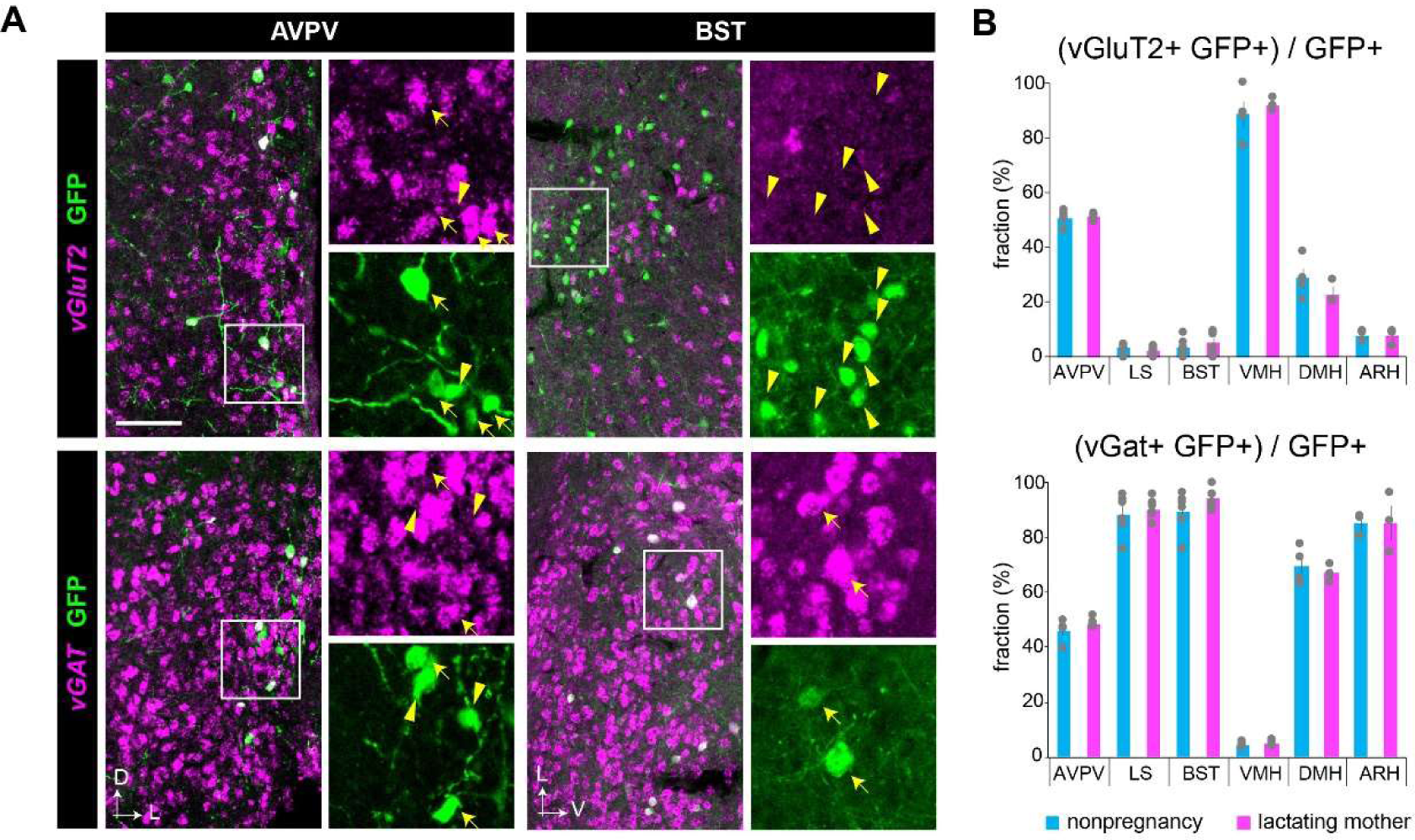
Cell-type analysis of rabies-labeled neurons. (A) We analyzed six brain regions (AVPV, LS, BST, VMH, DMH, and ARH), as these are the prominent presynaptic structures of OT neurons (Figure 3) and discrete structures that are easily identified in sections. Typical examples are shown for AVPV and BST coronal sections with *vGluT2* or *vGAT* ISH (red), along with anti-GFP immunostaining (green). Scale bar, 100 μm. High magnification images to the right of each image correspond to boxed areas in the low magnification images. Yellow arrows indicate markers and GFP double-labeled cells, whereas yellow arrowheads show examples of GFP-positive but marker-negative cells. D, dorsal, L, lateral, V, ventral. (B) Average fractions of GFP-positive cells that are *vGluT2*-positive (top) or *vGAT*-positive (bottom) in the indicated brain regions. Error bars represent the SEM. Individual data are shown in gray. *n* = 3–6 animals for each brain region.

**Figure S3.**
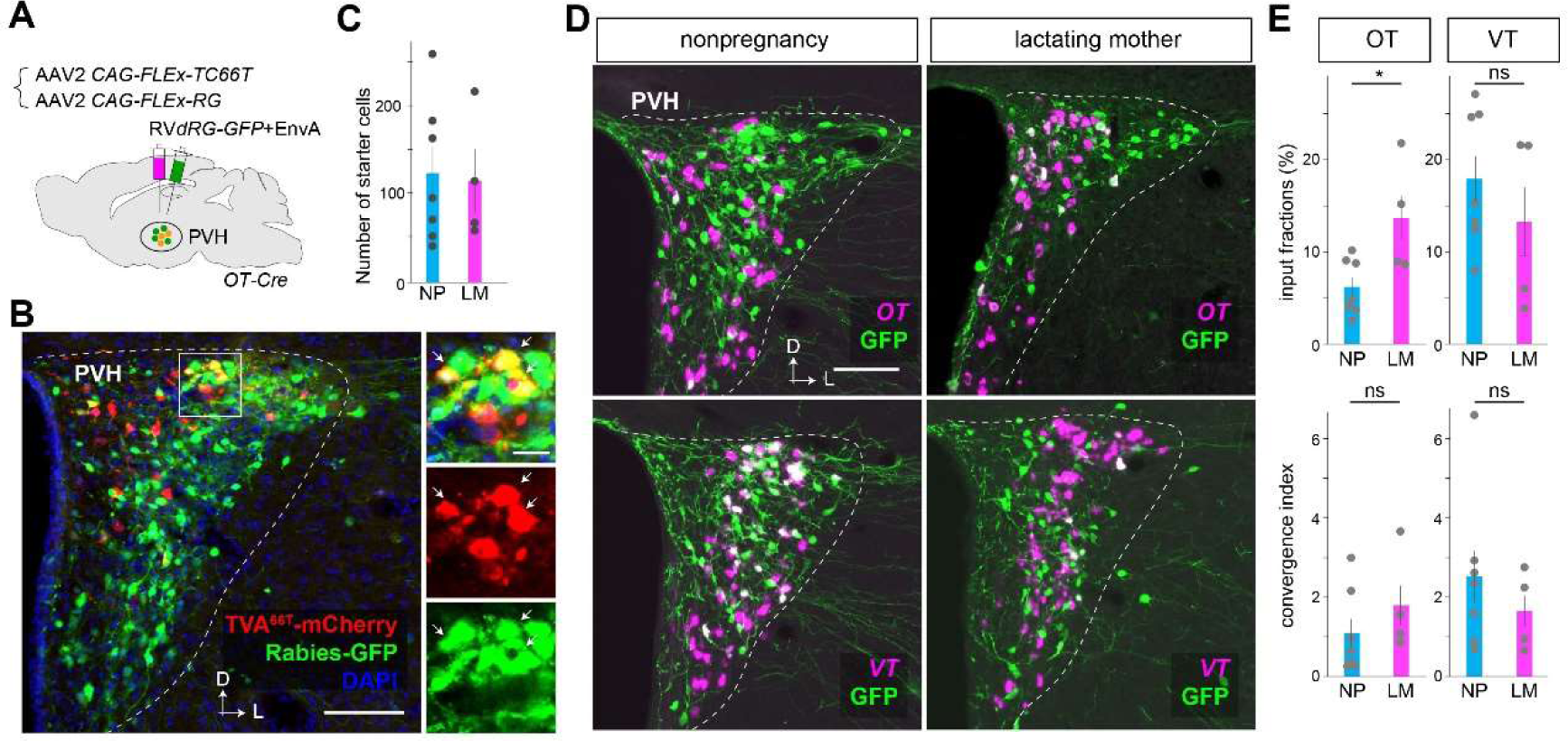
Local circuit input to OT neurons within the PVH. (A) Schematics of the experimental design. (B) Typical coronal section showing the starter cells that co-expressed TVA-mCherry (red) and rabies-GFP (green) in the PVH. D, dorsal, L, lateral. High magnification images to the right exhibiting red, green, and superimposed images correspond to the boxed area in the low magnification image. Arrows indicate some of the starter cells. Scale bar, 100 μm for low magnification images and 20 μm for high magnification images. (C) Quantification of starter cells per animal. No significant difference was found between nonpregnant (NP) and lactating mothers (LM) by the Wilcoxon rank-sum test (p = 0.6). *n* = 7 for NP and *n* = 3 for LM. (D) Representative coronal sections with *OT* or *vasotocin (VT)* ISH (magenta), along with anti-GFP immunostaining (green). Scale bar, 100 μm. D, dorsal, L, lateral. Although these two hormones were encoded by highly homologous genes, we carefully designed ISH probes to distinguish *OT* and *VT*. (E) Average input fraction (top) and convergence index (bottom) of *OT* and GFP dual-labeled cells (left) or *VT* and GFP dual-labeled cells (right). Error bars, SEM. Individual data are shown in gray. *, p = 0.023 by *t*-test, ns, p > 0.05 by two-sided *t*-test. Of note, the fraction and convergence index of OT-to-OT connections are underestimated because some presynaptic OT neurons may be recognized as starter cells. The estimated number of starter cells corresponded to 8.3% ± 2.4% and 13.4% ± 6.3% of OT+ neurons in NP and LM samples, respectively (p = 0.52 by the two-sided Wilcoxon rank-sum test). *n* = 7 for NP and *n* = 4 for LM.

**Figure S4.**
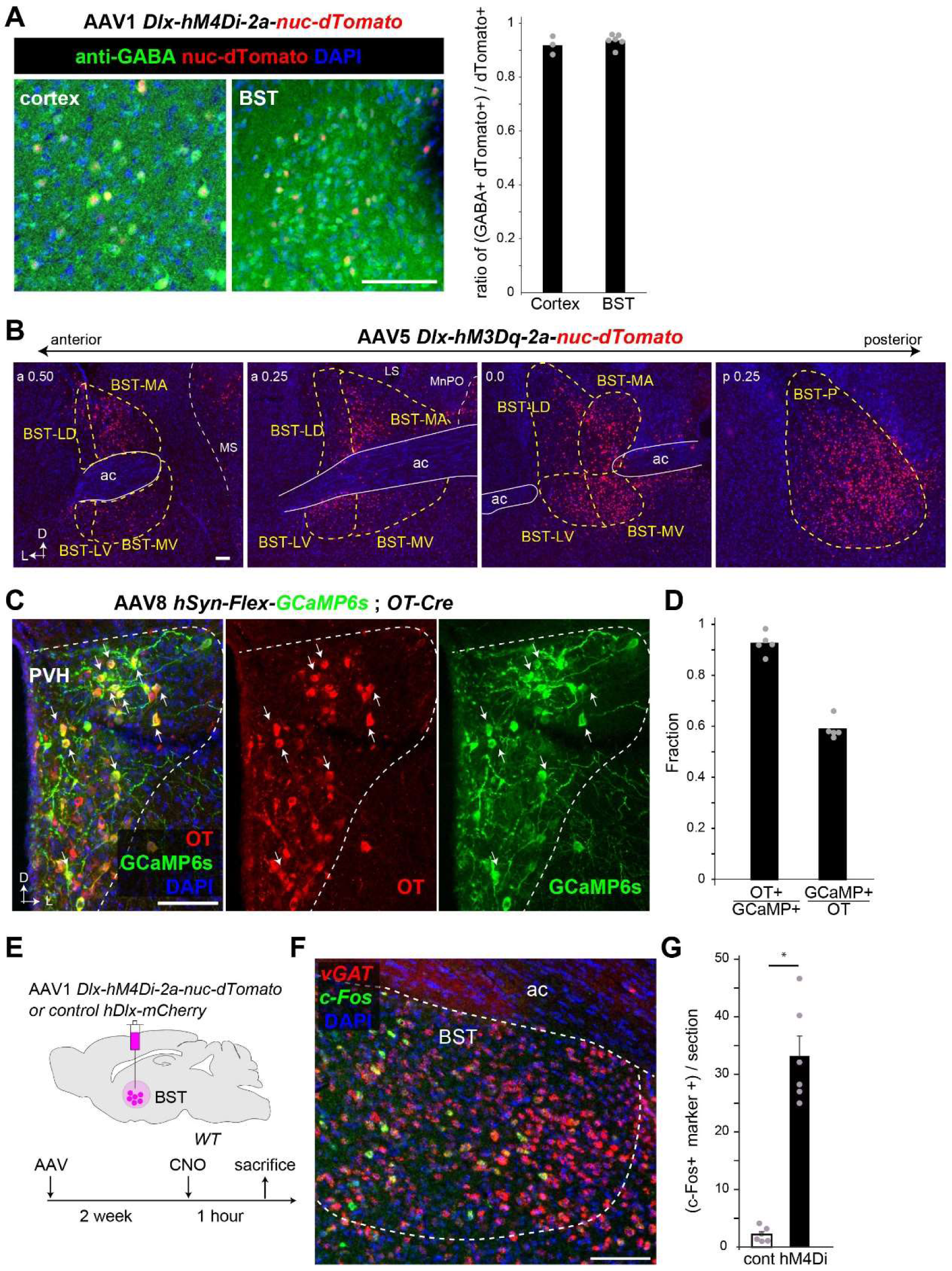
Additional data for histology supporting pharmacogenetic experiments, related to Figure 4. (A) Cell-type specificity of nuc-dTomato expression driven by the *Dlx* regulatory regions in the AAV vector. Brain slices of the cortex and BST were immunostained with anti-GABA antibody (green) and counterstained with DAPI (blue). The majority of dTomato+ were GABA+. *n* = 3 for the cortex and *n* = 6 for the BST. (B) Representative coronal sections showing the expression of hM3Dq inferred by the co-expressing nuc-dTomato marker in the BST and surrounding brain regions. The values in the top left indicate the location along the anterior–posterior axis (in mm) from the bregma. a, anterior, p, posterior. For abbreviations of brain regions, see the legend in Figure 4 and Table S2. (C) Typical example of a 30-μm coronal section of the PVH from *OT-Cre* female mice that had been injected with AAV9 *CAG-Flex-GCaMP6s* into the PVH. The section was stained with anti-OT (red) and anti-GFP (for GCaMP6s, green) antibodies and counterstained with DAPI (blue). Arrows indicate some GCaMP6s-positive cells. (D) Quantification of specificity (OT+/GCaMP6s+) and efficiency (GCaMP6s+/OT+). *n* = 5 mice each. Of note, the specificity and efficiency of GCaMP6s by the AAV-based method were comparable to those achieved by *OT-Cre; Ai162* (Figure 1B and 1C). (E) Because hM4Di-mediated inactivation of BST inhibitory neurons did not show any effect on the photometric peaks of OT neurons (Figure 4D, 4F, and 4G), we aimed to analyze whether this manipulation could impact the BST as assessed by *c-Fos* expression as a proxy of neural activation. Schematics of the experimental design and timeline are shown. At 1 h following saline or CNO injection, we obtained brain sections for ISH to detect *c-Fos* mRNA expression. (F) Typical example of a 30-μm coronal section of the PVH from hM4Di+ female mice stained with *vGAT* (a marker of inhibitory neurons shown in red) and *c-Fos* (shown in green) RNA probes. (G) Quantification of *c-Fos+ vGAT+* dual positive neurons in the BST per 30-μm coronal section. The hM4Di-group showed a significant increase of *c-Fos*+ inhibitory neurons in the BST. *, p < 0.05 by the Wilcoxon rank-sum test (*n* = 6). Due to technical reasons, we could not detect *c-Fos*+ cells simultaneously with dTomato epifluorescence, and therefore, we could not assess whether *c-Fos* induction happened in hM4Di+ cells. Silencing hM4Di+ inhibitory neurons by CNO might activate some local BST neurons via the disinhibition of local inhibitory circuits. Although the underlying mechanisms are unknown, these data imply that hM4Di in the BST can impact local neural activities. D, dorsal, L, lateral. Scale bars, 100 μm.

**Supplementary movie 1: Dynamics of OT neural activities during parturition, related to Figure 2.**

The *OT-Cre; Ai162* female mouse is about to give birth to her second pup in this parturition. Top, side (left), and bottom (right) views of the cage. Bottom, fiber photometry trace of GCaMP6s signals (the same sample shown in Figure 1F). The photometric peaks are indicated by arrowheads. The timing of abdominal contractions and pup delivery are represented by blue and red vertical dotted lines, respectively. Of note, abdominal contractions often occur 10–15 s after OT-PAs. The video is shown at 4× real-time speed.

**Supplementary movie 2: Photometric peak relative to suckling behaviors by pups, related to Figure 2.**

The *OT-Cre; Ai162* mother mouse returns to the nest to start breastfeeding three pups. Top, bottom view of the cage. Bottom, raster plots of behaviors of the mother and pups and a fiber photometry trace of GCaMP6s signals (the same sample shown in Figure 2F). In this sample, the photometric peak appears at about 4 min after the start of suckling by the third pup. The video is shown at 15× real-time speed.

## STAR☆METHODS

### KEY RESOURCES TABLE

| REAGENT or RESOURCE | SOURCE | IDENTIFIER |
| --- | --- | --- |
| **Antibodies** |  |  |
| horseradish peroxidase (HRP) conjugated anti-Dig antibody | Roche Applied Science | cat#11207733910; RRID:AB_514500 |
| rat anti-mCherry | Chromotek | cat#5f8; RRID:AB_2336064 |
| rabbit anti-OT | Immunostar | cat#20068; RRID:AB_572258 |
| donkey anti-rat IgG, Cy3 conjugated | Jackson Immuno Researchcat | #712-165-153; RRID:AB_2340667 |
| goat anti-chicken IgY, Alexa Fluor 488 | Jackson Immuno Research | cat#703-545-155; RRID:AB_2340375 |
| rabbit anti-GABA | Sigma-Aldrich | Cat# A2052, RRID:AB_477652 |
| chicken anti-GFP antibody | Aves Labs | cat#GFP-1020; RRID:AB_2307313 |
| donkey anti-rabbit IgG, Alexa Fluor 488 | Thermo Fisher Scientific | cat# A32790; RRID:AB_2762833 |
| donkey anti-rabbit IgG, Alexa Fluor 488 | Thermo Fisher Scientific | cat# A32794; RRID:AB_2762834 |
| horseradish peroxidase (HRP) conjugated anti-Flu antibody | Perlin Elmer | cat# NEF710001EA, RRID:AB_2737388 |
| **Bacterial and virus strains** |  |  |
| AAV serotype 5 *CAG-FLEX-TC<sup>b</sup>* | Miyamichi et al., 2013 (Custom made by UNC Vector Core) | N/A |
| AAV serotype 8 *CAG-FLEX-RG* | Miyamichi et al., 2013 (Custom made by UNC Vector Core) | N/A |
| AAV serotype 2 *CAG-FLEX-TC<sup>66T</sup>* | Miyamichi et al., 2013 (Custom made by UNC Vector Core) | N/A |
| AAV serotype 2 *CAG-FLEX-RG* | Miyamichi et al., 2013 (Custom made by UNC Vector Core) | N/A |
| AAV serotype 5 *hDlx-hM3Dq-nuc-dTomato* | Dimidschstein J. et al., 2016 (Custom made by Fukushima Medical Univ. by using plasmid Addgene #83897) | N/A |
| AAV serotype 1 *hDlx-hM4Di-nuc-dTomato* | Dimidschstein J. et al., 2016 (Custom made by Fukushima Medical Univ. by using plasmid Addgene #83896) | N/A |
| AAV serotype 1 *mDlx-ChR2-mCherry* | Dimidschstein J. et al., 2016 (Custom made by Canadian Neurophotonics Platform by using plasmid Addgene #83898) | N/A |
| AAV serotype 9 *CAG-FLEX-GCaMP6s* | Chen TW. et al., 2013 | RRID: Addgene_100842 |
| Rabies *dG-GFP+EnvA* | Reconstructed and amplified in the lab | Osakada and Callaway, 2013 |
| Experimental models: Cell lines |  |  |
| B7GG cell line | Gift from Dr. Edward Callaway | Osakada and Callaway, 2013 |
| BHK-EnvA cell line | Gift from Dr. Edward Callaway | Osakada and Callaway, 2013 |
| HEK293-TVA800 cell line | Gift from Dr. Edward Callaway | Osakada and Callaway, 2013 |
| Experimental models: Organisms/strains |  |  |
| Mouse: *OT-Cre* (Oxytocin-Ires Cre) | JAX Laboratories | Jax# 024234 |
| Mouse: *Ai162 (TIT2L-GC6s-ICL-tTA2)* | JAX Laboratories | Jax#031562 |
| Oligonucleotides |  |  |
| Primers for RNA probes, see **Histology and histochemistry** below | This paper (Custom made by Thermo Fisher Scientific) | N/A |
| Recombinant DNA |  |  |
| Plasmid: *pAAV CAG-FLEX-TC<sup>b</sup>* | Miyamichi et al., 2013 | RRID: Addgene_48332 |
| Plasmid: *pAAV CAG-FLEX-RG* | Miyamichi et al., 2013 | RRID: Addgene_48333 |
| Plasmid: *pAAV CAG-FLEX-TC<sup>66T</sup>* | Miyamichi et al., 2013 | RRID: Addgene_48331 |
| Plasmid: *pAAV-hDlx-GiDREADD-dTomato-Fishell-5* | Dimidschstein J. et al., 2016 | RRID: Addgene_83896 |
| Plasmid: *pAAV-hDlx-GqDREADD-dTomato-Fishell-4* | Dimidschstein J. et al., 2016 | RRID: Addgene_83897 |
| Plasmid: *pAAV-mDlx-ChR2-mCherry-Fishell-3* | Dimidschstein J. et al., 2016 | RRID: Addgene_83898 |
| mouse whole brain cDNA | GenoStaff | cat#MD01 |
| Software and algorithms |  |  |
| Doric Neuroscience Studio Software | Doric Lenses, Inc | 5.4.1.1 |
| MATLAB | Mathworks | R2020b Update 2 (9.9.0.1524771) |
| Other |  |  |

### LEAD CONTACT AND MATERIALS AVAILABILITY

Further information and requests for materials and data used in this study should be directed to and will be fulfilled by the Lead Contact, Kazunari Miyamichi, at. This study did not generate new unique reagents.

### EXPERIMENTAL MODEL AND SUBJECT DETAILS

Experimental protocols utilizing the rabies virus followed Biosafety Level 2 (P2/P2A) procedures approved by the biosafety committee of the RIKEN Center for Biosystems Dynamics Research (BDR). All animal procedures followed animal care guidelines approved by the Institutional Animal Care and Use Committee of the RIKEN Kobe branch. Wild-type C57BL/6j mice were purchased from Japan SLC (Shizuoka, Japan) for histological and mating experiments. *OT-Cre* (Jax#024234) and *Ai162* (*TIT2L-GC6s-ICL-tTA2*, Jax#031562) were purchased from the Jackson Laboratory. Animals were housed under a regular 12-h dark/light cycle with *ad libitum* access to food and water.

### METHOD DETAILS

#### Viral preparations

The following AAV vectors were generated by UNC viral core using the corresponding plasmids as described in the original literature^26^. The titer of the AAV was estimated by quantitative PCR methods and shown as genome particles (gp) per milliliter.

AAV serotype 5 *CAG-FLEx-TC^b^* (9.3 × 10^12^ gp/ml)

AAV serotype 8 *CAG-FLEx-RG* (2.8 × 10^12^ gp/ml)

AAV serotype 2 *CAG-FLEx-TC^66T^* (1.0 × 10^12^ gp/ml)

AAV serotype 2 *CAG-FLEx-RG* (1.3 × 10^12^ gp/ml)

The following AAV vectors were generated by the viral vector cores of Fukushima Medical University School of Medicine and Canadian Neurophotonics Platform by using the corresponding plasmids (Addgene #83896 and #83897) described in the original literature^28^.

AAV serotype 5 *hDlx-hM3Dq-nuc-dTomato* (1.0 × 10^13^ gp/ml)

AAV serotype 1 *hDlx-hM4Di-nuc-dTomato* (8.9 × 10^12^ gp/ml)

AAV serotype 1 *mDlx-ChR2-mCherry* (1.3 × 10^13^ gp/ml)

AAV serotype9 CAG-FLEx-GcaMP6s (1.7 × 10^13^ gp/ml), which was originally described in ref. 41, was obtained from Addgene (#100842).

Rabies *dG-GFP*+EnvA was prepared by using B7GG and BHK-EnvA cells (kindly gifted by Ed Callaway) according to the published protocol^42^. The EnvA-pseudotyped RV*dG-GFP*+EnvA titer was estimated to be 1.0 × 10^9^ infectious particles/ml based on serial dilutions of the virus stock, followed by infection of the HEK293-TVA800 cell line.

#### Stereotactic injection

For targeting AAV or rabies virus into a certain brain region, stereotactic coordinates were first defined for each brain region based on the Allen Brain Atlas^43^. Mice were anesthetized with 65 mg/kg ketamine (Daiichi-Sankyo) and 13 mg/kg xylazine (Sigma-Aldrich) via intraperitoneal injection and head-fixed to the stereotactic equipment (Narishige). For rabies tracing experiments (Figure 3 and Figure S2), 200 nl of a 1:1 mixture of AAV5 *CAG-FLEx-TC^b^* and AAV8 *CAG-FLEx-RG* was injected into the PVH at a speed of 50 nl/min using a UMP3 pump regulated by Micro-4 (World Precision Instruments). For local circuit mapping (Figure S3), 200 nl of a 1:1 mixture of AAV2 *CAG-FLEx-TC^66T^*and AAV2 *CAG-FLEx-RG* was injected into the PVH, followed by 200 nl of Rabies *dG-GFP*+EnvA injection to the same coordinate 2 weeks later. For pharmacogenetic experiments (Figure 4), 200 nl of AAV5 *hDlx-hM4Di-2a-nuc-dTomato*, AAV1 *hDlx-hM3Dq-2a-nuc-dTomato*, or AAV1 *mDlx-ChR2-mCherry* as a negative control, was injected into the BST. The following coordinates were used (distance in millimeters from the Bregma for the anterior [A]– posterior [P] and lateral [L] positions, and from the brain surface for the ventral [V] direction)]: BST, A 0.2, L 0.6, V 3.7; and PVH, A -0.75 L 0.2, V 4.5. After viral injection, the incision was sutured, and the animal was warmed using a heating pad to facilitate recovery from anesthesia. The animal was then returned to the home cage.

### Fiber photometry

For fiber photometry recording, *Ai162/+; OT-Cre/+* double heterozygous female mice were used. For some experiments in Figure 4, *OT-Cre* single heterozygous female mice that had been injected with AAV9 *CAG-FLEx-GCaMP6s* into the PVH were also used. A 400-μm core, 0.5 NA optical fiber (Thorlabs, cat#FP400URT) was implanted immediately above the PVH. After the surgery, animals were crossed with stud males and housed in the home cage until recording. We performed Ca^2+^ imaging by delivering excitation lights (465-nm modulated at 309.944 Hz and 405-nm modulated at 208.616 Hz) and collected emitted fluorescence by using the integrated Fluorescence Mini Cube (Doric, iFMC4_AE(405)_E(460-490)_F(500-550)_S). Light collection, filtering, and demodulation were performed using Doric photometry setup and Doric Neuroscience Studio Software (Doric Lenses, Inc.). The 405-nm signal was recorded as a background (non–calcium-dependent), and the 465-nm signal reported calcium-dependent GcaMP6s excitation/emission. The power output at the tip of the fiber was about 5 μW. The signals were initially acquired at 12 kHz and then decimated to 120 Hz for recording to disk. As the 405-nm signals were flat when 465-nm signals showed peaks in parturition (Figure 1F) or lactation (not shown), we did not apply further background subtraction methods. We used a 2-Hz low-pass filter before the analysis of the photometric peaks of OT neurons.

For the analyses, we used a homemade MATLAB code. Briefly, the ΔF/F was calculated by 100 × (F_*t*_ – F_0_)/F_0_, where F_*t*_ was the recorded signal at time=*t* and F_0_ was the average of signals in the whole recording period. Because the height of peaks in each mother varied considerably, as the optical fiber location relative to the PVH was variable, to identify the photometric peaks reliably, we first selected several visually obvious peaks to estimate the peak height of that animal. The photometric peaks of OT neurons were then automatically detected by using the findpeaks function in MATLAB, with the peak threshold of half of the estimated peak height, and the FWHM threshold over 4 s. To show the peri-event traces of the peaks (Figures 1G, 2B, and 2I), we extracted the ΔF/F data from –10 s to +15 s around the peak (the local maximum point of ΔF/F). We then adjusted the median fluorescence of the –8 to –3 s baseline period to zero to align multiple data.

Videos from a side infrared camera (DMK33UX273; The Imaging Source) were synchronized with fiber photometry acquisition. The camera from the bottom (QWatch; IODATA) was matched with the side camera using the mouse motion and light conditions. For the analysis of parturition, based on the bottom view of the video record, the delivery timing was defined as the moment when the entire pup was outside the vagina. We extracted the ΔF/F data from –10 min to +5 min around each delivery and analyzed only the peaks detected before delivery because those after delivery might be included in the next delivery event. For the heat map representation (Figure 1H), ΔF/F was normalized to the maximum value within that animal in a –10 min to +5 min period of the analysis. To compare peak height (Figure 2J, 2K, and 2N and Figure S1J), we calculated the averaged peak height of one condition normalized to the averaged peak height of another condition of the same animal to obtain the normalized ΔF/F. For the analysis of photometric peaks during lactation, PPD1 was defined as 1 day after the day of parturition. In the cross-fostering experiments (Figure 2L–2N), we used the postnatal day 0–1 pup as ‘young’ and the postnatal day 10–16 pup as ‘elder’.

To monitor the sucking by pups during lactation (Figure 2F and 2G), hairs of the abdomen of PPD9 mothers were removed under slight anesthesia. At PPD10, the mother and three pups were placed in an acryl cage with nest materials and a minimum amount of woody chips. One hour after connecting to the photometry system, we carefully placed two cameras (Qwatch, IODATA) just beneath the nest from the bottom. We quantified the latency to the first photometric peak from the time when the mother returned to the nest or the time when pups started suckling behaviors.

To assess the crouching duration of mothers (Figure 4H), we measured the amount of time when mothers stayed in their nest and interacted with pups within a 1-h time window starting from 30 min after saline or CNO injection.

#### Pharmacogenetics

Neural activation or inactivation experiments were performed at least 2 weeks after the injection of AAV driving hM3Dq or hM4Di. Then, 0.3 ml of 400 μg/ml CNO (Sigma-Aldrich cat#C0832) dissolved in saline or 0.3 ml of saline was intraperitoneally injected into the animal during the fiber photometry imaging sessions.

#### Histology and histochemistry

For the quantitative analysis of the trans-synaptic tracing samples and the histochemical analyses, the experimental mice were anesthetized with a lethal amount of sodium pentobarbital, sacrificed, and perfused with phosphate-buffered saline (PBS) followed by 4% paraformaldehyde (PFA) in PBS. Brain tissues were post-fixed with 4% PFA in PBS overnight at 4 °C, cryoprotected with 30% sucrose solution in PBS at 4 °C for 24– 48 h, and embedded in the O.C.T. compound (Tissue-Tek, cat#4583). We collected 30-μm coronal sections of the whole brain using Cryostat (model#CM1860; Leica) and placed them on MAS-coated glass slides (Matsunami). Unless otherwise noted, every third (Figures 3 and 4 and Figure S2 and S4) or fifth (Figure 1 and Figure S3) coronal brain section was analyzed for quantification, and compensated data (×3 or ×5) were represented.

To map the long-distance input to OT neurons (Figure 3), every third of the coronal brain section was imaged using a slide scanner (Zeiss Axio Scan.Z1) with a 20× objective lens (NA 0.8). Image processing was performed semi-automatically using ImageJ macro (National Institutes of Health, Bethesda, MD). Briefly, regions of interest (ROIs) were manually set for each brain region by using the DAPI channel (showing brain structures but not GFP+ cells) based on the Allen Brain Atlas^43^ by annotators who were blinded to the experimental conditions. For each ROI, the raw fluorescent image for GFP or mCherry was processed using open and median filters. The labeled cells were detected by using the “Threshold” and “Analyze Particles” commands. The starter cells were defined by those detected as cells using both GFP and mCherry channels. In Figure 3E, the *d′* value was calculated as (μ_*mother*_ – μ_*nonpregnancy*_) divided by (σ_*mother*_ + σ_*nonpregnancy*_), where μ and σ denote the average and standard deviation, respectively. A d′ value larger than 0 means that OT neurons in lactating mothers receive more input.

Next, 30-μm coronal sections containing the target brain region were subjected to ISH (Figures S2–S4), as described previously^44^. To generate cRNA probes, DNA templates were amplified by PCR from the C57BL/6j mouse genome or whole-brain cDNA (Genostaff, cat#MD-01). T3 RNA polymerase recognition site (5’-AATTAACCCTCACTAAAGGG) was added to the 3’ end of the reverse primers. Primer sets to generate DNA templates for cRNA probes are as follows (the first one, forward primer, the second one, reverse primer): *OT* 5’- TGTGCTGGACCTGGATATGCG; 5’-CGCCGTGCACAATCCGAATC *VT* 5’- GAATGAAGGGAGTCGAGGGTT; 5’-TCCCCACCCCAGAAATAGAGAC *vGluT2-1* 5’-TAGCTTCCTCTGTCCGTGGT; 5’-GGGCCAAAATCCTTTGTTTT *vGluT2-2* 5’-CCACCAAATCTTACGGTGCT; 5’-GGAGCATACCCCTCCCTTTA *vGluT2-3* 5’-CTCCCCCATTCACTACCTGA; 5’-GGTCAGGAGTGGTTTGCATT *vGAT1-1* 5’-CCTGGTCTGGACAGCATCTC; 5’-GCTATGGCCACATACGAGTC *vGAT1-2* 5’-GTCAATGTGGCGCAGATCAT; 5’-CCTAGTCCTCTGCGTTGGTT *cFos-1* 5’-AGCGAGCAACTGAGAAGACTG; 5’-ATCTCCTCTGGGAAGCCAAG *cFos-2* 5’-CCAGTCAAGAGCATCAGCAA; 5’-CATTCAGACCACCTCGACAA

DNA templates (500–1000 ng) amplified by PCR were subjected to *in vitro* transcription with DIG (cat#11277073910)- or Flu (cat#11685619910)-RNA labeling mix and T3 RNA polymerase (cat#11031163001) according to the manufacturer’s instructions (Roche Applied Science). When possible, two or three independent RNA probes for the same gene were mixed to increase the signal/noise ratio.

For ISH combined with anti-GFP staining, after hybridization and washing, sections were incubated with horseradish peroxidase (HRP)-conjugated anti-Dig (Roche Applied Science cat#11207733910, 1:500) and anti-GFP (Aves Labs cat#GFP-1020, 1:500) antibodies overnight. Signals were amplified by TSA-plus Cyanine 3 (AKOYA Bioscience, NEL744001KT, 1:70 in 1× plus amplification diluent) for 25 min, followed by washing, and then GFP-positive cells were visualized by anti-chicken Alexa Fluor 488 (Jackson Immuno Research cat#703-545-155, 1:250). PBS containing 50 ng/ml 4’,6-diamidino-2-phenylindole dihydrochloride (DAPI; Sigma-Aldrich, cat#D8417) was used for counter nuclear staining.

For dual-color ISH (Figure S4E–S4G), DIG-labeled *vGAT* probes and Flu-labeled *c-Fos* probes were mixed for hybridization. DIG-positive cells were visualized with TSA-plus Cyanine 3 (AKOYA Bioscience, NEL744001KT, 1:70 in 1× plus amplification diluent), and Flu-positive cells were detected with anti-Flu antibody (PerkinElmer, NEF710001EA, 1:250 in blocking buffer) followed by TSA-plus biotin (AKOYA Bioscience, NEL749A001KT, 1:70 in 1× plus amplification diluent) and streptavidin-Alexa Fluor 488 (Life Technologies, 1:250).

For immunohistochemistry, sections were washed three times with PBS containing 0.3% Tween-20 (PBST) for 10 min and treated with 5% normal donkey serum (NDS; Southern Biotech, cat#0030-01) in PBST for 1 h at room temperature for blocking. The following primary antibodies were used in this study: rat anti-mCherry (Chromotek cat#5f8, 1:500–1:1000), chicken anti-GFP (Aves Labs cat#GFP-1020, 1:500), rabbit anti-OT (Immunostar cat#20068, 1:1000), and rabbit anti-GABA (Sigma-Aldrich cat#A2052, 1:1000). These antibodies were diluted into 5% NDS in PBST for 3 h at room temperature or overnight at 4 °C. Signal-positive cells were detected by the following secondary antibodies: anti-rat Cy3 (Jackson Immuno Research cat#712-165-153, 1:250), anti-chicken Alexa Fluor 488 (Jackson Immuno Research cat#703-545-155, 1:250), anti-rabbit Alexa Fluor 488 (Thermo Fisher Scientific cat#A32790, 1:250), and anti-rabbit Alexa Fluor 555 (Thermo Fisher Scientific cat#A32794, 1:250) diluted into PBST for 2 h at room temperature or overnight at 4 °C. Sections were washed once with PBST for 10 min, treated with PBS containing DAPI for 20 min, rinsed with PBS, and mounted with cover glass using Fluoromount (Diagnostic BioSystems cat#K024).

To analyze the efficiency and specificity of GCaMP6s expression (Figure 1C and Figure S4C), 30-μm coronal sections containing PVH were stained with anti-OT and anti-GFP (for GCaMP6s) antibodies. Then, OT+, GCaMP6s+, and dual-positive cells were manually counted from at least five coronal sections of the PVH per animal. For the *c-Fos* assay (Figure S4E–S4G), AAV5 *hDlx-hM4Di-2a-nuc-dTomato* was injected into the BST of 8 week-old wild-type C57BL/6j female mice. After the surgery, animals were singly housed for 2 weeks. They were sacrificed 1 h after saline or CNO injection, and brain samples were processed for dual-color ISH to detect *c-Fos* and *vGAT* mRNA expression. Dual-labeled cells were manually counted for at least five coronal sections of the BST per animal. To analyze the distribution of hM3Dq+ cells in the BST subdivisions (Figure 4J and 4K, Figure S4B), based on the shape of the anterior commissure^45^, we annotated labeled cells into one of five subdivisions: medial anterior, medial ventral, lateral dorsal, lateral ventral, and posterior. Of note, the medial division posterior part, medial division posterior intermediate part, and lateral division posterior part in the atlas^45^ were collectively grouped as “posterior” in this work.

Sections were imaged using an Olympus BX53 microscope with a 4× (NA 0.16) or 10× (NA 0.4) objective lens equipped with a cooled CCD camera (DP80; Olympus) or Zeiss Axio Scan.Z1 with a 20× (NA 0.8) objective lens. Images were processed in ImageJ and Photoshop CC (Adobe).

## Authors’ contributions

H.Y. and K.M. designed the study. H.Y. performed the fiber photometry recordings, retrograde trans-synaptic tracing, and pharmacogenetic manipulation experiments with technical support from M.H., K.T., and C.H.L. S.K. and K.K. provided the AAV *Dlx-DREADD* viruses. H.Y. and K.M. wrote the manuscript with assistance from all co-authors.

## Acknowledgments

We wish to thank K. Inada, T. Goto, and K. Murata for technical advice on fiber photometry, S. Irie for technical support, E. Callaway for sharing the B7GG, BHK-EnvA, and HEK293-TVA800 cell lines, and all the laboratory members for their help and critical reading of the manuscript. This work was supported by Kakenhi 17K14948, 20K15907, and RIKEN diversity promotion grants to H.Y., AMED Brain/MINDS program 20dm0207052h0004 to K.K., and JST PREST program JPMJPR1789, Kakenhi 18H02548 and 20K20589, a Takeda Science Foundation Research Grant, and The Uehara Memorial Foundation Research Grant to K.M.

## References

1. Armstrong, W. E. Central Nervous System Control of Oxytocin Secretion during Lactation. Physiology of Reproduction 5th *edition*, 527-560 (2015).

2. Brunton, P. J. R. J. A. Maternal Brain Adaptations in Pregnancy. Physiology of Reproduction 5th *edition*, 1957-2026 (2015).

3. Froemke, R. C. C. I. Oxytocin and Brain Plasticity. Principles of Gender-Specific Medicine, 161–182 (2017).

4. Wakerley, J. B. & Lincoln, D. W. The milk-ejection reflex of the rat: a 20- to 40-fold acceleration in the firing of paraventricular neurones during oxytocin release. J Endocrinol 57, 477–493, doi:10.1677/joe.0.0570477 (1973).

5. Lincoln, D. W. & Wakerley, J. B. Factors governing the periodic activation of supraoptic and paraventricular neurosecretory cells during suckling in the rat. J Physiol 250, 443–461, doi:10.1113/jphysiol.1975.sp011064 (1975).

6. Sutherland, R. C., Juss, T. S. & Wakerley, J. B. Prolonged electrical stimulation of the nipples evokes intermittent milk ejection in the anaesthetised lactating rat. Exp Brain Res 66, 29–34, doi:10.1007/BF00236198 (1987).

7. Belin, V. & Moos, F. Paired recordings from supraoptic and paraventricular oxytocin cells in suckled rats: recruitment and synchronization. J Physiol 377, 369–390, doi:10.1113/jphysiol.1986.sp016192 (1986).

8. O’Byrne, K. T., Ring, J. P. & Summerlee, A. J. Plasma oxytocin and oxytocin neurone activity during delivery in rabbits. J Physiol 370, 501–513, doi:10.1113/jphysiol.1986.sp015947 (1986).

9. Paisley, A. C. & Summerlee, A. J. Activity of putative oxytocin neurones during reflex milk ejection in conscious rabbits. J Physiol 347, 465–478, doi:10.1113/jphysiol.1984.sp015076 (1984).

10. Summerlee, A. J. & Lincoln, D. W. Electrophysiological recordings from oxytocinergic neurones during suckling in the unanaesthetized lactating rat. J Endocrinol 90, 255–265, doi:10.1677/joe.0.0900255 (1981).

11. Lewis, E. M. et al. Parallel Social Information Processing Circuits Are Differentially Impacted in Autism. Neuron 108, 659–675 e656, doi:10.1016/j.neuron.2020.10.002 (2020).

12. Zhang, B. et al. Reconstruction of the Hypothalamo-Neurohypophysial System and Functional Dissection of Magnocellular Oxytocin Neurons in the Brain. Neuron 109, 331–346 e337, doi:10.1016/j.neuron.2020.10.032 (2021).

13. Luo, L., Callaway, E. M. & Svoboda, K. Genetic Dissection of Neural Circuits: A Decade of Progress. Neuron 98, 256–281, doi:10.1016/j.neuron.2018.03.040 (2018).

14. Hung, L. W. et al. Gating of social reward by oxytocin in the ventral tegmental area. Science 357, 1406–1411, doi:10.1126/science.aan4994 (2017).

15. Tang, Y. et al. Social touch promotes interfemale communication via activation of parvocellular oxytocin neurons. Nat Neurosci 23, 1125–1137, doi:10.1038/s41593-020-0674-y (2020).

16. Marlin, B. J., Mitre, M., D’Amour J, A., Chao, M. V. & Froemke, R. C. Oxytocin enables maternal behaviour by balancing cortical inhibition. Nature 520, 499–504, doi:10.1038/nature14402 (2015).

17. Knobloch, H. S. et al. Evoked axonal oxytocin release in the central amygdala attenuates fear response. Neuron 73, 553–566, doi:10.1016/j.neuron.2011.11.030 (2012).

18. Carcea, I. et al. Oxytocin neurons enable social transmission of maternal behaviour. Nature, doi:10.1038/s41586-021-03814-7 (2021).

19. Gunaydin, L. A. et al. Natural neural projection dynamics underlying social behavior. Cell 157, 1535–1551, doi:10.1016/j.cell.2014.05.017 (2014).

20. Wu, Z. et al. An obligate role of oxytocin neurons in diet induced energy expenditure. PLoS One 7, e45167, doi:10.1371/journal.pone.0045167 (2012).

21. Daigle, T. L. et al. A Suite of Transgenic Driver and Reporter Mouse Lines with Enhanced Brain-Cell-Type Targeting and Functionality. Cell 174, 465–480 e422, doi:10.1016/j.cell.2018.06.035 (2018).

22. Dubois-Dauphin, M., Armstrong, W. E., Tribollet, E. & Dreifuss, J. J. Somatosensory systems and the milk-ejection reflex in the rat. II. The effects of lesions in the ventroposterior thalamic complex, dorsal columns and lateral cervical nucleus-dorsolateral funiculus. Neuroscience 15, 1131–1140, doi:10.1016/0306-4522(85)90257-x (1985).

23. Honda, K. & Higuchi, T. Effects of unilateral electrolytic lesion of the dorsomedial nucleus of the hypothalamus on milk-ejection reflex in the rat. J Reprod Dev 56, 98–102, doi:10.1262/jrd.09-090e (2010).

24. Lebrun, C. J., Poulain, D. A. & Theodosis, D. T. The role of the septum in the control of the milk ejection reflex in the rat: effects of lesions and electrical stimulation. J Physiol 339, 17–31, doi:10.1113/jphysiol.1983.sp014699 (1983).

25. Miyamichi, K. et al. Cortical representations of olfactory input by trans-synaptic tracing. Nature 472, 191–196, doi:10.1038/nature09714 (2011).

26. Miyamichi, K. et al. Dissecting local circuits: parvalbumin interneurons underlie broad feedback control of olfactory bulb output. Neuron 80, 1232–1245, doi:10.1016/j.neuron.2013.08.027 (2013).

27. Theofanopoulou, C., Gedman, G., Cahill, J. A., Boeckx, C. & Jarvis, E. D. Universal nomenclature for oxytocin-vasotocin ligand and receptor families. Nature 592, 747–755, doi:10.1038/s41586-020-03040-7 (2021).

28. Dimidschstein, J. et al. A viral strategy for targeting and manipulating interneurons across vertebrate species. Nat Neurosci 19, 1743–1749, doi:10.1038/nn.4430 (2016).

29. Armbruster, B. N., Li, X., Pausch, M. H., Herlitze, S. & Roth, B. L. Evolving the lock to fit the key to create a family of G protein-coupled receptors potently activated by an inert ligand. Proc Natl Acad Sci U S A 104, 5163–5168, doi:10.1073/pnas.0700293104 (2007).

30. Valtcheva, S. et al. Neural circuitry for maternal oxytocin release induced by infant cries. bioRxiv, 2021.2003.2025.436883, doi:10.1101/2021.03.25.436883 (2021).

31. Summerlee, A. J. Extracellular recordings from oxytocin neurones during the expulsive phase of birth in unanaesthetized rats. J Physiol 321, 1–9, doi:10.1113/jphysiol.1981.sp013967 (1981).

32. Thirtamara Rajamani, K., et al. Efficiency of cell-type specific and generic promoters in transducing oxytocin neurons and monitoring their neural activity during lactation. Sci Rep 11, 22541, doi:10.1038/s41598-021-01818-x (2021).

33. Son, S. et al. Wiring diagram of the oxytocin system in the mouse brain. bioRxiv, 2020.2010.2001.320978, doi:10.1101/2020.10.01.320978 (2020).

34. Freda, S. N. et al. Brainwide input-output architecture of paraventricular oxytocin and vasopressin neurons. bioRxiv, 2022.2001.2017.476652, doi:10.1101/2022.01.17.476652 (2022).

35. Inada, K. et al. Plasticity of neural connections underlying oxytocin-mediated parental behaviors of male mice. Neuron, doi:10.1016/j.neuron.2022.03.033 (2022).

36. Theodosis, D. T., Piet, R., Poulain, D. A. & Oliet, S. H. Neuronal, glial and synaptic remodeling in the adult hypothalamus: functional consequences and role of cell surface and extracellular matrix adhesion molecules. Neurochem Int 45, 491–501, doi:10.1016/j.neuint.2003.11.003 (2004).

37. Moos, F. et al. Oxytocin in the bed nucleus of the stria terminalis and lateral septum facilitates bursting of hypothalamic oxytocin neurons in suckled rats. J Neuroendocrinol 3, 163–171, doi:10.1111/j.1365-2826.1991.tb00259.x (1991).

38. Dulac, C. & Wagner, S. Genetic analysis of brain circuits underlying pheromone signaling. Annu Rev Genet 40, 449–467, doi:10.1146/annurev.genet.39.073003.093937 (2006).

39. Kim, S. R. & Kim, S. Y. Functional Dissection of Glutamatergic and GABAergic Neurons in the Bed Nucleus of the Stria Terminalis. Mol Cells 44, 63–67, doi:10.14348/molcells.2021.0006 (2021).

40. Dewey, K. G. Maternal and fetal stress are associated with impaired lactogenesis in humans. J Nutr 131, 3012S–3015S, doi:10.1093/jn/131.11.3012S (2001).

41. Chen, T. W. et al. Ultrasensitive fluorescent proteins for imaging neuronal activity. Nature 499, 295–300, doi:10.1038/nature12354 (2013).

42. Osakada, F. & Callaway, E. M. Design and generation of recombinant rabies virus vectors. Nat Protoc 8, 1583–1601, doi:10.1038/nprot.2013.094 (2013).

43. Lein, E. S. et al. Genome-wide atlas of gene expression in the adult mouse brain. Nature 445, 168–176, doi:10.1038/nature05453 (2007).

44. Ishii, K. K. et al. A Labeled-Line Neural Circuit for Pheromone-Mediated Sexual Behaviors in Mice. Neuron 95, 123–137 e128, doi:10.1016/j.neuron.2017.05.038 (2017).

45. Franklin, K. B. J. & Paxinos, G. Paxinos and Franklin’s The mouse brain in stereotaxic coordinates. Fourth edition. edn, (Academic Press, an imprint of Elsevier, 2013).

